# Comprehensive Proteomic, Ultrastructural and Functional Analysis of Cryopreservation Effects on Marine Mussel Oocytes

**DOI:** 10.1101/2024.11.25.625176

**Authors:** Sofía Blanco, Sara Campos, Patricia Reboreda, Estefanía Paredes, Angel P. Diz

## Abstract

Cryopreservation is a valuable tool for preserving marine genetic diversity and supporting selective breeding in aquaculture. The blue mussel (*Mytilus galloprovincialis*) is an important species for the aquaculture industry and a useful model for studying cryopreservation in marine invertebrates. Declining mussel seed availability over the past decade, combined with the growing demand for year-round hatchery production, has increased interest in developing effective cryopreservation methods for gametes and embryos. However, cryopreservation remains challenging, particularly for marine invertebrate oocytes, as it requires balancing cryoprotectant cytotoxicity with sufficient cell protection during freezing and thawing. Moreover, the molecular, structural, and functional changes associated with oocyte cryopreservation in invertebrates are still poorly understood. This study examines the effects of two commonly used cryoprotectants, dimethyl sulfoxide (DMSO) and ethylene glycol (EG), on *M. galloprovincialis* oocytes under a monitored slow freezing (MSF) protocol. Using an integrative analytical approach combining proteomics, functional assessments, and ultrastructural analyses, we provide a comprehensive understanding of cryoprotectant-induced changes. Our findings reveal that cryoprotectants cause significant proteomic alterations, with DMSO having more pronounced effects. These alterations are associated with increased oxidative stress, an insufficient antioxidant response, and disruptions in meiosis restart mechanisms, ultimately delaying larval development. Cryoprotectant-induced oxidative stress may also weaken the oocyte membrane, leading to rupture during the MSF protocol and subsequent fertilisation failure post-thawing. These results provide evidence that EG is less cytotoxic than DMSO and suggest that supplementing cryoprotectants with antioxidants and membrane stabilizers could enhance success rates in the cryopreservation of oocytes from mussels and other marine invertebrate species.

## 1. INTRODUCTION

The cryopreservation of gametes is relevant in both conservation biology and aquaculture, serving as a long-term genetic reservoir [31,39]. In aquaculture, this approach preserves genetic diversity, enhances selective breeding programs and facilitates the production of hybrids to exploit heterotic effects. It also enables on-demand offspring production, independent of natural breeding seasons, and extends the reproductive lifespan of valuable specimens, mitigating genetic loss from unforeseen events or environmental changes. This technology ensures a stable supply of high-quality broodstock, essential for meeting the growing demand for seafood while maintaining ecological balance.

Despite its potential, the cryopreservation of fish and shellfish gametes has seen limited progress and generally poor outcomes, particularly with oocytes [17]. Factors such as low membrane permeability and large yolk content in oocytes may hinder the penetration of cryoprotectant agents (CPAs), leading to intracellular ice formation and significant cellular damage [1,13]. Dimethyl sulfoxide (DMSO) and ethylene glycol (EG) are two of the most commonly used CPAs, widely applied to minimise ice formation in cryopreserved sperm, oocytes, and embryos across both aquatic and non-aquatic species [2,3,5,10,30,33,42,54,62,67,71]. Oocyte structural and compositional characteristics may contribute to their sensitivity to the chilling process; however, the exact reasons for their poor cryopreservation outcomes remain unclear. Another factor to consider is the broad variation among species, which may lead to differential cytotoxic effects of CPAs or varying sensitivity to freezing-thawing protocols [8,11].

Many marine invertebrate species, including several bivalve molluscs, are of great interest to the aquaculture industry. A common feature among them is the external fertilisation, which facilitates the induction, collection, management, and cryopreservation of large quantities of gametes. Blue mussels (*Mytilus* spp.) serve as excellent model organisms in reproductive and developmental biology studies of bivalve species with high interest for aquaculture. The increasing demand for mussel seed, coupled with a notable drop in seed availability in some regions over the last decade [4,47], highlights the need for alternatives beyond natural supply. Cryopreservation has emerged as an alternative method for sustainable mussel seed production, addressing seasonal limitations while ensuring the continuous availability of genetically diverse and resilient mussel populations. This approach benefits commercial aquaculture and contributes to ecological conservation by minimising disturbance to marine beds and rocky shores during seed collection.

However, the success rates of sperm cryopreservation in blue mussels (*Mytilus galloprovincialis*) have been limited and heterogeneous [15,35]. The same applies to other marine invertebrates [34,50]. Challenges with oocyte cryopreservation have been even more pronounced in terms of fertilisation and subsequent successful larval development, with low to nil success rate across various bivalve species, including blue mussels (*M. galloprovincialis*), green-lipped mussels (*Perna canaliculus*), or Pacific oysters (*Crassostrea gigas*) [2,23,33,60,62]. Similar problems have been reported in other marine invertebrates, such as sea urchins and starfish, where low post-thaw fertilisation rates have been observed [26,49]. On the other hand, evaluations of CPA-induced damage in mussel gametes have also been limited, often relying on viability assays and oxidative stress markers [23,62]. These methods, along with techniques for estimating DNA damage, and enzymatic assays, have been used to evaluate the alterations caused by the freezing process itself in marine gametes [31]. However, the results from these analyses are often limited to molecular targets hypothesised to be involved in cryodamage based on prior knowledge.

High-throughput molecular techniques focussed on gene expression analysis have been less frequently used, despite their utility in both discovery and hypothesis-driven studies. These techniques can help fill current knowledge gaps about the effects of CPAs and cryodamage on gametes. While transcriptomic (RNA-seq) analyses in mammalian cryopreserved gametes are more common [24,28,36,70], such studies are scarce for marine organisms and have focused exclusively on cryopreserved sperm [16,63], with no existing studies in marine invertebrates. Furthermore, proteomics has emerged as a powerful and highly informative approach because proteins represent the phenotype at molecular level and vary in response to environment changes or external stimuli [19,40]. This underscores the value of proteomics in comprehensively understanding cytotoxicity and cryodamage on gametes, through the exploration of the intricate effects on accessory and reproductive proteins, which work all together to ensure fertilisation [22]. Similar to RNA-seq studies, proteomics has been applied to mammalian cryopreserved gametes [6,29,38,64], but studies on marine organisms are also scarce and again only on sperm [69,75], with no data on marine invertebrates. Furthermore, despite the robustness offered by studying cryodamage from a holistic and multidisciplinary perspective, multi-technique approaches that integrate functional, structural, and molecular analyses have not yet been conducted on cryopreserved gametes from marine organisms.

In this study, we analysed changes in the oocyte proteome of *Mytilus galloprovincialis* induced by two commonly used CPAs (DMSO and EG) and a monitored slow freezing (MSF) technique. These findings were integrated with results from ultrastructural (electron microscopy) and functional analyses, including assessments of fertilisation and development rates.

## 2. METHODS

### 2.1. Mussel sampling and spawning induction

About one hundred of specimens from marine mussels *M. galloprovincialis* (4-5 cm shell length) were collected in the rocky shore of Toralla island, Vigo (NW coast of Spain, 42°12’06.7“N 8°47’57.5“W) during the reproductive season (March 2021), transported to the marine station (ECIMAT, University of Vigo) and maintained in seawater that was filtered through a 1 µm pore size and treated with UV light (hereafter referred to as treated seawater), with a temperature maintained at 13±1°C and a salinity of 34.5±1.1‰. Spawning was induced following a thermal shock procedure [52], oocyte samples were gathered using a Pasteur pipette and washed thrice through a 41 µm mesh with treated seawater to remove debris. The integrity of oocytes was inspected through optical microscopy, and only oocyte samples with good quality (i.e., exhibiting rounded shape, homogeneous colour and intact membrane) were retained for further analysis. This procedure was repeated until oocyte samples were collected from seven different females, hereafter referred to as biological replicates.

### 2.2. Application of cryoprotectant agents and monitored slow freezing method

Each sample of oocytes obtained from four biological replicates was divided into five sub-samples, each containing approximately 40 oocytes/mL, that were designated for the application of the two different cryoprotectant agents (CPAs), i.e., dimethyl sulfoxide (DMSO) and ethylene glycol (EG), either with or without the subsequent application of a monitored slow-freezing technique (hereafter treatments); together with control oocytes as outlined in **Table 1**. For the CPA treatment, oocytes underwent a 15-minute incubation period in either 1.5 M DMSO or 1.5 M EG solution prepared in treated seawater at room temperature (19±1°C). The CPA concentration and incubation time selected were those used in previous studies with *M. galloprovincialis* oocytes [33,62]. Following this, either a flash frozen using N_2_(l) or a monitored slow freezing (MSF) method was applied to each subsample treated with the same CPA. The MSF procedure involved a cooling ramp consisting of conditioning the cells at 4°C for 2 minutes, cooling from 4°C to −10°C at a rate of 1°C/min, holding at −10°C for 2 minutes for manual seeding to induce slow and stable crystal nucleation [32] and cooling from - 10°C to −35°C (−1°C/min) in a programmable Controlled Freezer (Cryologic Lpt. Australia). Once −35°C was reached, samples were plunged and stored in N_2_(l) [2]. Control samples consisted on a subsample of oocytes from the same biological replicates after 15 min incubation in treated seawater to rule out the incubation time as a potential confounding factor. For the subsequence proteomics analysis (section 2.5), both control and CPA-treated samples were stored in N_2_(l). On the other hand, to verify the *in vitro* fertilisation and to D-larvae development rate of cryopreserved oocytes with the MSF treatment (DMSO - MSF and EG - MSF), cryopreserved oocytes were thawed using the method described by Adams *et al.* [2]. Subsequently, they were fertilised and screened following the protocol detailed in section 2.3.

**Table 1.** Sample Codes and Treatments for Oocyte Analysis. . Control samples: Oocytes were incubated for 15 minutes in treated seawater (i.e., filtered through a 1 µm pore size and treated with UV light). DMSO and EG-treated samples: Oocytes were incubated in either 1.5 M DMSO or 1.5 M EG for 15 minutes. Cryopreserved samples using the monitored slow freezing method (DMSO-MSF and EG-MSF): Oocytes were treated with either DMSO or EG and then subjected to monitored slow freezing (MSF) for 35 minutes. All samples, including the control and the CPA-treated ones, were snap-frozen and stored in liquid nitrogen (N_2_(l)) before proteomic analysis.

| Code | Treatment |
| --- | --- |
| Control | treated seawater (15') |
| DMSO | 1.5 M DMSO in treated seawater (15') |
| EG | 1.5 M EG in treated seawater (15') |
| DMSO-MSF | 1.5 M DMSO in treated seawater (15') + monitored slow freezing (35') + $N_2(l)$ |
| EG-MSF | 1.5 M EG in treated seawater (15') + monitored slow freezing (35') + $N_2(l)$ |

### 2.3. *In vitro* fertilisation and early embryo development analysis

To determine if CPAs impact fertilisation and early embryonic development rate, oocyte subsamples at a concentration of 40 oocytes/mL from a new set of three biological replicates (see section 2.1) were treated with both CPAs following the procedure described in the section 2.2 and subsequently fertilised at room temperature while maintaining an approximate 10:1 sperm-to-oocyte ratio. The sperm, obtained from a single male, showed optimal motility after optical microscopy inspection. The fertilisation rate was assessed 90 minutes’ post-treatment, categorising oocytes as fertilised if they displayed the release of the first/second polar body, or were in the early two-cell embryo stage. Subsequently, the early development rate was assessed by incubating the fertilised eggs at 18°C for 72h and evaluating their development into D-larvae. Data were collected after fixing the samples with 0.2% formalin to assess the number of normally developed D-larvae, as well as undeveloped oocytes (whether fertilised or not) and other earlier developmental stages ranging from 16-cell embryos to trochophores (hereafter “embryos”), which precede the D-larvae stage. Each biological replicate and treatment included three technical replicates. Counts were conducted for at least 85 subjects per technical replicate, covering unfertilised oocytes, fertilised oocytes, embryos, or D-larvae, using an inverted microscope. To assess whether there were significant differences in fertilisation and early development rates between control and treatment samples, a one-way ANOVA with repeated measures (p<0.05) was performed. An arcsine transformation was applied to the average fertilisation and D-larvae rates, which were calculated from technical replicates within each biological replicate. For post-hoc comparisons, Dunnett’s multiple comparisons test was performed. All statistical analyses were done with GraphPad Prism vs8.0.1.

### 2.4. Scanning electron microscope analysis of oocyte ultrastructure

To detect alterations in the oocyte’s ultrastructure using scanning electron microscopy (SEM), an aliquot of either control or treated (e.g., exposed to a specific CPA and cryopreserved) oocyte samples were fixed in 1% glutaraldehyde in treated seawater. They were washed three times in 0.1M cacodylate buffer (pH 7.2) and postfixed in 1% osmium tetroxide in the same buffer for 2 hours. After three additional washes in the same buffer, they were subjected to dehydration using an increasing concentration of acetone in water (30, 50, 70, 80, 90 and 100%). Samples were then subjected to drying via the critical point technique using the *Baltec CPD 020* critical point desiccator, and were subsequently attached to a carbon adhesive support and coated with gold for observation in the *JEOL JSM6700F* field emission SEM. Finally, photographs were taken at different magnifications. To estimate the rate of damaged oocytes, we captured representative panoramic images at 250X magnification. Subsequently, detailed photographs were taken of the most abundant oocyte types in the sample at higher magnifications, where individual oocytes were visible at 1,300X to 1,600X, and their surface details were observed at 10,000X.

### 2.5. Protein extraction, purification and trypsin digestion

The total protein content was extracted from each of twenty oocyte samples (4 biological replicates per treatment; see section 2.2 and **Table 1**) and solubilised in a homogenisation buffer (7 M Urea, 2 M Thiourea, and 4% CHAPS), assisted by sonication (Branson Digital Sonifier 250) operating at 20% amplitude with 5 blasts (5 s each, with 10-s pauses) on ice. After centrifugation for 20 min at 16000g, the pellet was discarded, and the protein supernatant stored at −80°C until further use. Protein concentration was quantified using the Bradford method with bovine serum albumin (BSA) as a standard, and adjusted to 0.5 μg/μl in LC-MS grade water. Subsequently, proteins were purified through the acetone precipitation method, followed by reduction using tris(2-carboxyethyl) phosphine (TCEP) and alkylation using iodoacetamide (IAA), and digested with trypsin prior to isobaric labelling (further details are provided in Diz and Sánchez-Marín [20]).

### 2.6. Isobaric peptide labelling and LC-MS/MS analysis

Tryptic peptides derived from the different samples and treatments were labelled with different isobaric tandem mass tags from the TMT10plex Kit^TM^ following the manufacturer’s instructions as illustrated in **File S1–worksheet(ws)A**. Subsequently, multiplexed peptides underwent pre-fractionation using the Pierce High pH Reversed-Phase Peptide Fractionation Kit^TM^ to enhance the analytical resolution of the mass spectrometry analysis. The resulting peptides from the eight fractions were dried using a vacuum concentrator and reconstituted in 0.5% formic acid for tandem mass spectrometry analysis (MS/MS). The MS/MS analysis of peptides from each fraction was performed using a LQT-Orbitrap ELITE coupled to a Proxeon EASY-nLC 100 UHPLC system (Thermo Fisher). Peptides were separated on an RP column (EASY-Spray, 50 cm × 75 μm ID, PepMap C18, 2 μm particles, 100 Å pore size) with a 10 mm precolumn (Accucore XL C18). The gradient elution used 0.1% formic acid (mobile phase A) and acetonitrile with 0.1% formic acid (mobile phase B) over 240 minutes with a linear gradient from 5% to 35% B at a flow rate of 300 nL/min. Ionization occurred in a nanosource at 1.95 kV with a capillary temperature of 275 °C, and the Orbitrap set at a mass resolving power of 120,000. Peptide analysis was conducted in positive mode (1 μscan; 400–1600 amu), followed by 10 data-dependent HCD MS/MS scans (1μscans). The normalized collision energy was set at 38%, with an isolation width of 1.5 amu. Dynamic exclusion was activated with a repeat count of 1, a repeat duration of 30 s, an exclusion duration of 80 s, and a relative exclusion width of 10 ppm. Unassigned charged ions were excluded from the analytical process.

### 2.7. Protein identification and quantification

RAW files containing mass spectrometry data were used for protein identification and quantification analysis using the PEAKS® X Pro software (Bioinformatics Solutions Inc., Waterloo, Canada). Experimental MS and MS/MS spectra were compared with theoretical spectra generated from proteins present in a customised database containing 23,990 protein predictions. These predictions were derived from unpublished RNA-Seq data from our laboratory obtained from mature *M. galloprovincialis* female gonads (**File S2**). Protein predictions were annotated using a threshold *e*-value of 1 × 10^−3^ against Swiss-Prot database (March 1, 2023; 610,265 sequences). A complementary annotation against the nrNCBI protein database, restricted to Mollusca taxa (January 18, 2023; 1,457,555 sequences), was also conducted in order to refine the annotation of those proteins showing either non-informative protein descriptions or no annotation against Swiss-Prot. The customised database also included a set of common contaminant (https://www.thegpm.org/crap) and decoy sequences to calculate the false discovery rate (FDR). Protein identifications were carried out by allowing a maximum of two trypsin missed cleavages, and precursor and fragment ion mass tolerance values were set at 10 ppm and 0.02 Da, respectively. The search was conducted allowing also for two fixed post-translational modifications (TMT10plex labelling and carbamidomethylation on cysteine residues), in addition to more than 300 PTMs as variable modifications while also considering potential non-synonymous nucleotide mutations. Only proteins identified with at least one unique peptide (quality score ≥ 10), an FDR ≤ 1%, and signal from the reporter TMT ions were included in the quantitative analysis.

### 2.8. Differential protein abundance analysis

The protein abundances for each multiplexed sample in the two different TMT10plex experiments were normalised within and between experiments using a methodology inspired by CONSTANd [61], as described in Diz and Sánchez-Marín [20]. Only proteins with less than 50% of missing data were retained for analysis, and the remaining missing data were imputed by the minimum abundance value of the entire dataset. The normalised data was transformed to logarithmic scale (log2) to meet the normality and homoscedasticity assumptions required for parametric statistical analysis. Differential protein abundance analyses were performed with paired *t*-test between the control and the CPA-treated samples and between the latter and the cryopreserved ones. *P*-values were corrected for the multiple-hypothesis testing problem [18,58]. Proteins were considered to exhibit a significant differential abundance (whether up- or down-modulated relative to the control group) if they had a *q*-value < 0.1 and a SFisher test value lower than 0.05. In addition, a binomial exact test (p < 0.05) was performed to determine whether the number of up- and down-regulated proteins fits the theoretical ratio of 1:1. Protein functional analyses were carried out on significant modulated proteins with an effect size (fold change: FC) higher than 1.5 or lower than 0.67 for either up or down-regulation, respectively. We first performed an enrichment analysis of Kyoto Encyclopedia of Genes and Genomes (KEGG) molecular pathways using the Fisher exact test (FDR < 0.05). This analysis was conducted in OmicsBox v2.2, following enzyme annotation (*e*-value < 1×10^-3^) of all protein predictions derived from the reference transcriptome obtained from mature female gonads of *M. galloprovincialis* (**File S2**). The entire set of annotated proteins was used as the reference set. In addition, physical and functional protein-protein interaction networks were constructed among the modulated proteins using the STRING algorithm [59], relying on the proteome information available for the marine invertebrate *Strongylocentrotus purpuratus*. This species was selected due to its close phylogenetic proximity to our species among all the available options in the STRING database. Using STRING, we also performed an enrichment analysis (Fisher exact test, FDR < 0.05) of the GO (Gene Ontology) terms for the three main categories (BP: Biological Process, MF: Molecular Function, and CC: Cellular Component). Proteins for which interactions were detected were used as the test set, while the complete proteome of the selected organism (*S. purpuratus*) was used as the reference set. After performing the enrichment analysis, the GO terms with the highest specificity (more specialised “child” terms) involving at least 20 proteins were shown in the interaction network.

## 3. RESULTS

### 3.1. Effects of cryopreservation on fertilisation and early embryo development

While control oocytes from four females (F1 to F4) exhibited a good fertilisation and D-larvae transformation rate of 94.5% (±3.1%) and 82.1% (±7.2%) respectively, the corresponding thawed cryopreserved oocytes (DMSO-MSF and EG-MSF) showed no fertilisation (**File S1-wsB**). In this initial assessment, it remains unclear whether fertilisation failure can be attributed to the application of monitored slow freezing (MSF) and subsequent oocyte thawing procedure, or to the toxic effects of the cryoprotectant agents (CPAs). Consequently, the same test was repeated using oocyte samples from another three females (F5 to F7), which were treated with CPAs, either DMSO or EG, and their respective control samples (see section 2.3). In this second test, although no significant decrease in fertilisation rate was detected between control and CPA-treated oocytes (ANOVA, p = 0.23), a significant decrease in the rate of early development was observed in treated compared to control samples (**File S1-wsC** and **Figure 1**). This was detected through a decrease in the rate of D-larvae in CPA-treated samples compared to controls (p = 0.010) accompanied by a significant increase (p = 0.0097) in the rate of embryos that did not reach D-larvae stage. Indeed, while control oocytes developed to D-larvae in 93.2% (±5.0) of cases, the addition of either DMSO or EG resulted in a drop to 52.4% (±5.0) and 67.7% (±4.4), respectively. The rest of measured cells were mostly in earlier stages. While there was a slight trend toward higher fertilisation and D-larvae development rates (with increases of 8% and 15%, respectively) in the EG-treated oocytes compared to those treated with DMSO, these differences were not statistically significant. Finally, when assessing the rate of undeveloped oocytes, which included both unfertilised and fertilised oocytes that did not develop further and represented less than 10% in all cases, the DMSO treatment showed a significantly higher number of undeveloped oocytes compared to both the control (p = 0.013) and EG-treated (p = 0.020) samples.

**Figure 1.**
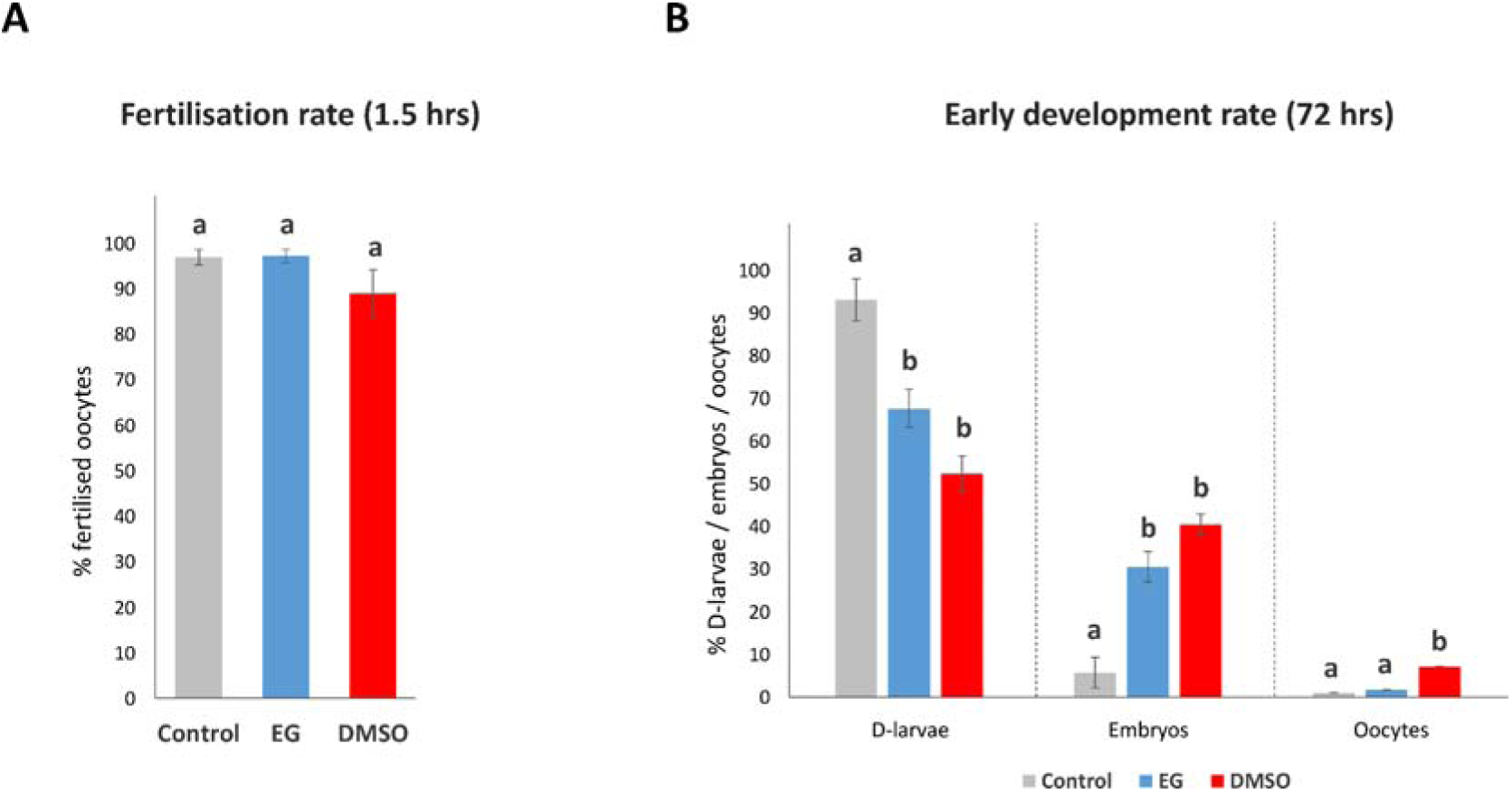
Fertilisation and Early Development Rates in Oocyte Samples. **(A)** Fertilisation percentage (mean ± SD; n=3) of control, EG, and DMSO-treated oocytes measured 1.5 hours after sperm addition, assessed by the first/second polar body release, the formation of the fertilisation membrane or, exceptionally, the visualisation of a two-cell embryo. **(B)** Early development percentages (mean ± SD; n=3) after 72 hours (post-fertilisation), comparing control oocytes with those treated with either DMSO or EG, specifically for D-shaped larvae (D-larvae), earlier embryological developmental stages (Embryos), and fertilised or unfertilised oocytes (Oocytes). More details about the experimental design and data collection are provided in section 2.3. Statistical significance (p < 0.05), indicated by different letters on the graph, reflects results from 1-way ANOVA post hoc tests, highlighting significant differences in developmental outcomes between treatments.

### 3.2 Effects of cryopreservation on oocyte ultrastructure

Investigating the viability of oocytes post-cryopreservation demands a meticulous examination. Initial observations under an optical microscope revealed apparently healthy, rounded, and intact oocytes in both control and cryopreserved samples following the thawing process. However, higher resolution scanning electron microscopy (SEM) allows for a comprehensive analysis aimed at uncovering ultrastructural defects within post-thawed cryopreserved oocytes that could explain the observed failure of fertilisation. Oocytes treated with DMSO or EG exhibited the same appearance as those in the control group, with 100% of the oocytes having an average diameter of 50 µm, without cell rupture and with an undamaged vitelline membrane (VM). However, the DMSO-MSF and EG-MSF samples showed 100% and 89% of damaged oocytes, respectively. These damages were evidenced by the outflow of intracellular contents or breaches in the oocyte envelope of variable depth. Notably, the 11% of oocytes without evidence of rupture in EG-MSF-treated samples did not exhibit the filamentary structures seen in the undamaged oocyte envelope (control and CPA-treated oocytes). **Figure 2** displays representative images at different magnifications of control oocytes, oocytes after incubation with either DMSO or EG, and oocytes treated with monitored slow freezing (MSF) followed by thawing (DMSO-MSF and EG-MSF).

**Figure 2.**
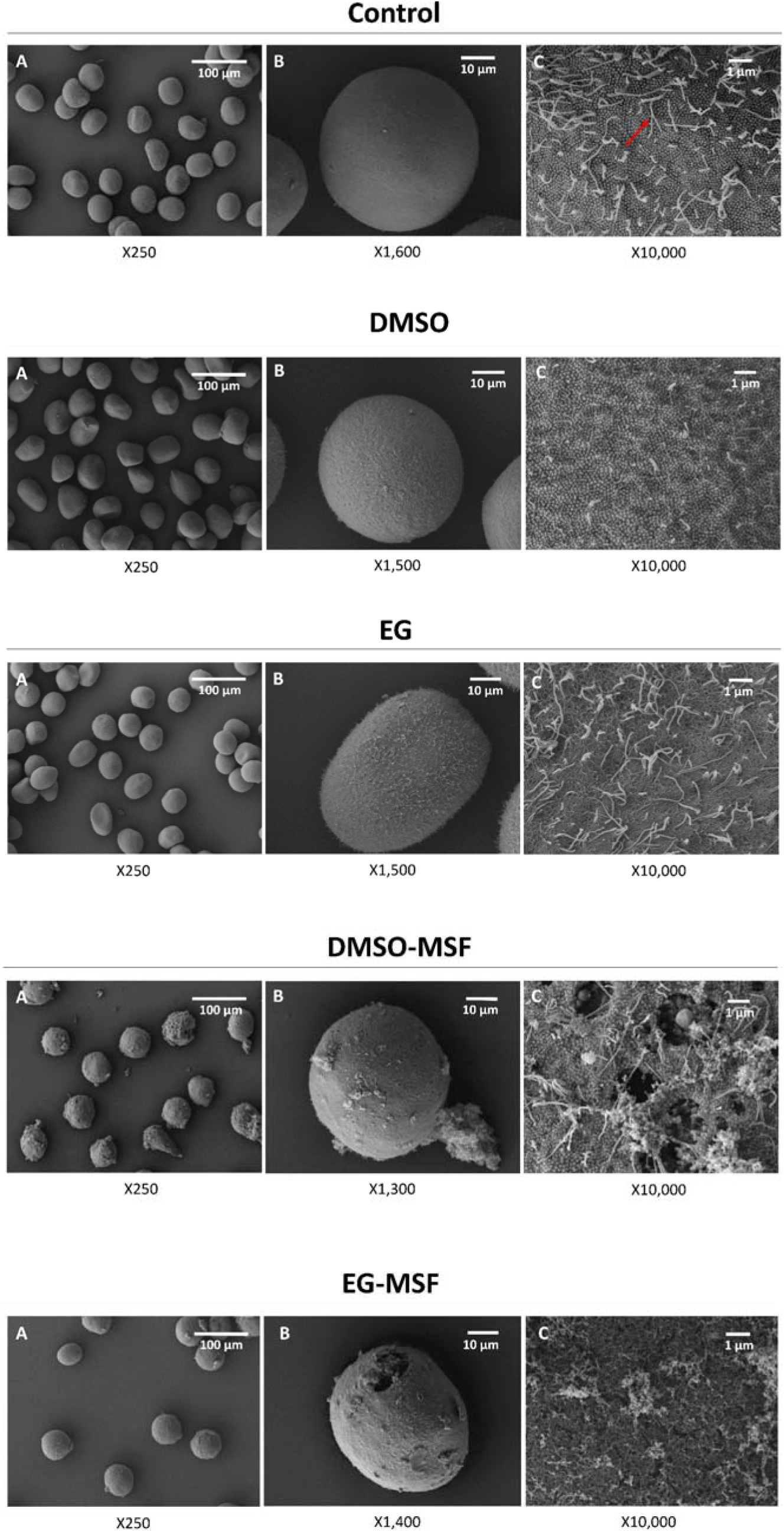
Ultrastructure Photographs of Control and Cryopreserved Oocytes. Ultrastructure photographs of control oocytes, oocytes treated with cryoprotectants (DMSO and EG), and cryoprotected oocytes subjected to monitored slow freezing (MSF) followed by thawing (DMSO-MSF and EG-MSF) were obtained using scanning electron microscopy (SEM). For each sample type, panoramic photographs (250X) show a representative fraction of the sample (**panel A**). Additionally, photographs at 1,300-1,600X magnification provide a representative example of the most abundant oocyte type in the sample (**panel B**). Detailed photographs at 10,000X (**panel C**) show the surface of the vitelline membrane (VM). In both control samples and samples treated exclusively with DMSO or EG, all oocytes appeared intact. However, in the DMSO-MSF-treated samples, 100% of the oocytes showed ruptures leading to leakage of cellular content. Similarly, in 89% of the EG-MSF-treated oocytes, ruptures were observed (**panel B**), while the remaining 11% showed no ruptures but exhibited a loss of the filamentous structures in the VM seen in control (red arrow, **panel C**) and CPA-treated oocytes.

### 3.3. Proteome alterations induced by cryoprotectant agents

A total of 40,315 peptides and 4,727 protein groups, hereafter referred to as ‘proteins’, have been identified and quantified in the analysed oocytes (**File S1-wsD and E).** Notably, the proteome analysis uncovered significant effects of cryoprotectant agents (CPAs), indicating a substantial modulation in the abundance of several proteins. Specifically, 1,207 (25.5%) exhibited significant changes in oocytes treated with DMSO, whereas 745 proteins (15.8%) showed alterations in those treated with EG (**File S1-wsF**). Furthermore, the number of up-regulated proteins was significantly higher than the number of down-regulated proteins, both in EG treatment (seven times more; exact binomial test, p < 0.001) and in DMSO treatment (three times more; exact binomial test, p < 0.001), with the latter inducing the most substantial size effects (**Figure 3**). Notably, the axonemal dynein heavy chain 5 protein exhibited an approximately 30-fold lower abundance in the DMSO treatment compared to control. However, the abundance of this protein was similar in EG-treated and control oocytes (FC = 1.17).

**Figure 3.**
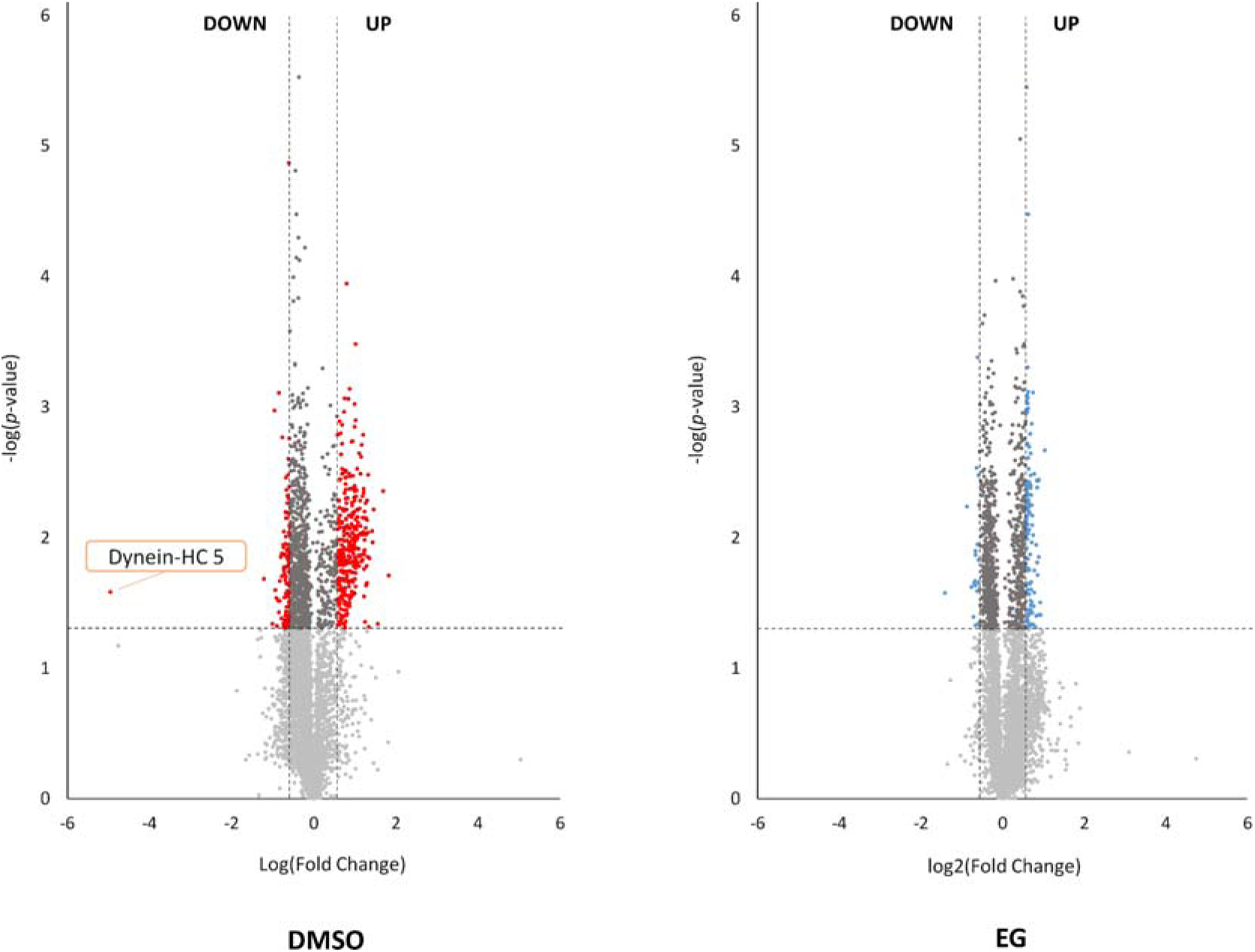
Volcano Plots of Protein Modulation in DMSO and EG Treated Oocytes. Volcano plots comparing DMSO and EG treated samples with control oocytes. In both cases, proteins with no significant modulation are shown in light grey, while the modulated ones (p < 0.05) with effect sizes (fold change) lower than 1.5 (whether up- or down-regulated) are in dark grey. Those modulated proteins exceeding this fold change threshold are highlighted in red for the DMSO treatment and blue for the EG treatment. The plots illustrate that DMSO treatment resulted in a greater number of modulated proteins, mostly up-rather than down-regulated, compared to EG treatment. Notice that fold change values in the DMSO treatment exhibit more extreme values. Notably, dynein-HC 5 displays an exceptionally lower relative abundance, with a 30-fold decrease in the DMSO-treated samples compared to controls.

#### 3.3.1. Proteins modulated with both cryoprotectant agents

Among the proteins that exhibited significant differential abundance (*q*< 0.01) in response to either DMSO or EG exposure, 496 were found to be common to both treatments, showing a remarkably high correlation in their effect sizes (Pearson’s correlation, *r* = 0.96, p < 0.001) (**File S1-wsG)**. Focussed on the 83 proteins displaying the highest effect sizes in both treatments (FC > 1.50 or < 0.67) we investigated the protein-protein interaction network. From that list, 58 proteins constituted a molecular network enriched in Biological Processes (BP) associated with the metabolism of carboxylic acid and organonitrogen compounds such as lipids and amino acids. Moreover, these proteins present mostly oxidoreductase and nucleotide-binding functions (Molecular Functions, MF), being mainly located in the mitochondrial matrix (Cellular Components, CC) (**Figure 4A**). All but two were up-regulated with both CPAs (see the results of GO terms enrichment analyses in **File S1-wsH,** and in **File S1-wsI** the list of proteins involved in the terms highlighted in **Figure 4A**). Notably, the identification of the oxidative stress biomarker superoxide dismutase (SOD) Fe-Mn among the mitochondrial oxidoreductases stands out. Also, the identification of mitochondrial heat shock proteins (Hsp) 70, 75, 10, and 60, which play pivotal roles in nucleotide binding and DNA repair mechanisms, along with the protease ClpP involved in the quality control system of mitochondrial proteins, is noteworthy.

**Figure 4.**
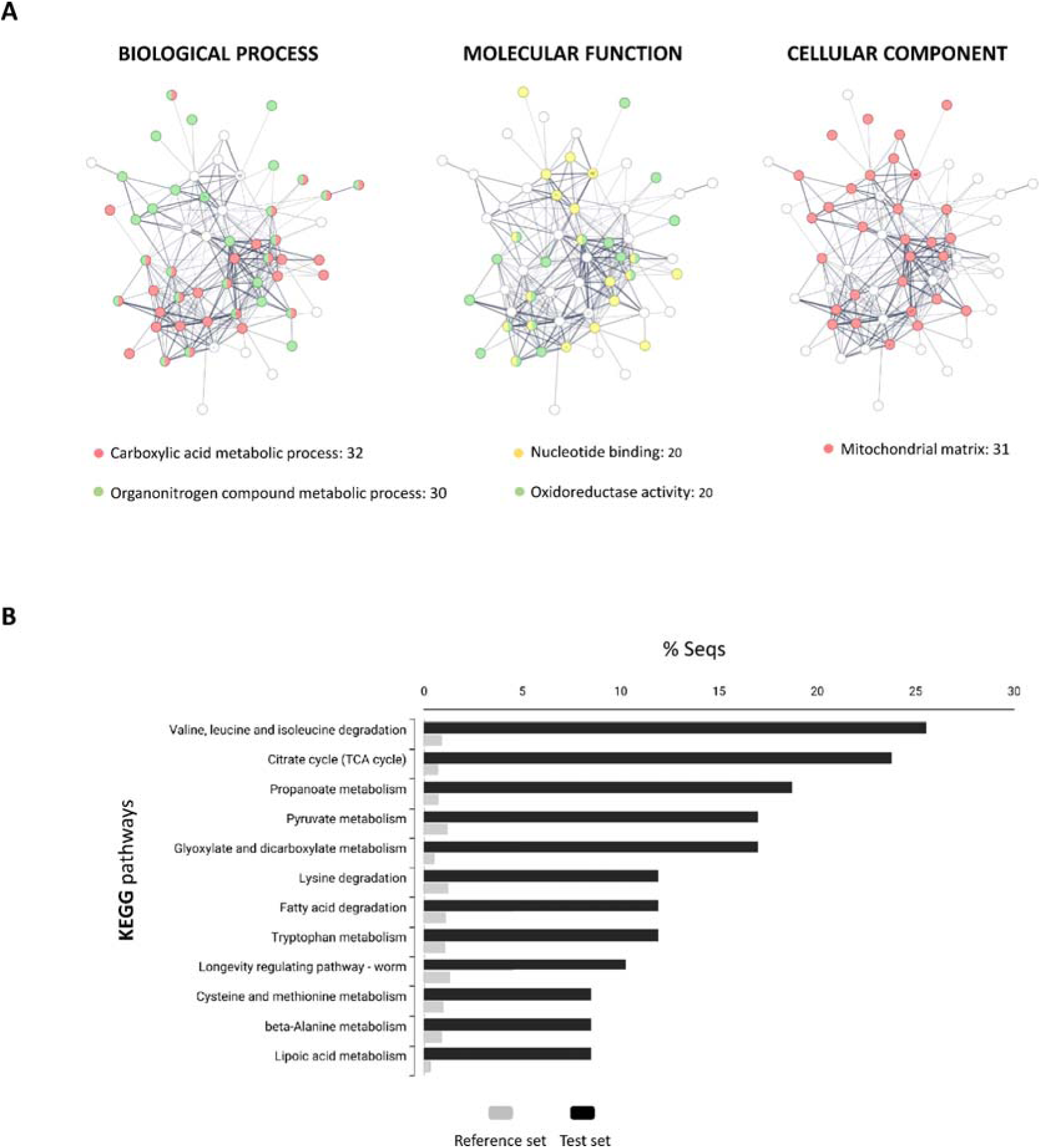
Interaction Networks and KEGG Molecular Pathways of Modulated Proteins. **A)** Physical and functional interaction networks were obtained through the analysis of 83 proteins modulated by both CPAs, with a q-value < 0.1 and fold changes greater than 1.5 for up-regulation or less than 0.67 for down-regulation. The most specific GO terms for Biological Process, Molecular Function, and Cellular Component categories that are enriched (Fisher exact test, FDR < 0.05) and involve more than 20 proteins in the network are shown. Next to each GO term, colour-coded in the interaction network, the number of proteins involved is indicated (a list of which can be found in **File S3-ws I**). **B)** KEGG molecular pathways that are significantly over-represented (Fisher exact test, FDR < 0.05) among the 83 proteins modulated by both CPAs (test set) compared to the total number of quantified proteins (reference set).

The results of KEGG pathways enrichment analysis using the same test set of 83 proteins is also in line with the up-regulation of lipid, amino acid and carboxylic acid (via the TCA cycle) metabolism (**Figure 4B**). The results did not vary substantially regardless of whether the entire protein database or only those proteins identified and quantified as a reference set were used. Also noteworthy is the finding that nine up-regulated enzymes were linked to the glyoxilate cycle and that four proteins were involved in the lipoic acid metabolism, a process described as a regulator of mitochondrial redox balance [57]. Beyond these metabolic changes, there is an overrepresentation of up-regulated enzymes linked to pathways associated with longevity regulation. These include the Hsp 60 and 70, the SOD Fe/Mn and the ATP-dependent protease ClpP, which play key roles in responding to stress and activating the mitochondrial unfolding protein response, a cellular mechanism controlling the misfolding of mitochondrial proteins under cellular stress conditions. For the detailed list of proteins involved in each pathway and their respective expression fold changes, please refer to **File S1-wsJ**.

#### 3.3.2. Distinctive protein modulations resulting from DMSO or EG exposure

Exclusively with DMSO treatment, we observed a significantly higher number of up-regulated proteins (320) compared to down-regulated proteins (100), indicating a threefold higher ratio of up to down-regulation (p < 0.001 in a binomial test against a 1:1 ratio). The upregulated proteins consistently exhibited enrichment in GO terms, as discussed in section 3.3.1. Furthermore, they unveiled associations with biological processes (BP) such as mitochondrial gene expression and sulfur metabolism, and molecular functions (MF) that included the purine ribonucleoside triphosphate binding (**File S1-wsK**). The involvement of sulfur metabolism was further supported by the KEGG pathways enrichment results (**File S3-wsL**). These pathways encompass sulphur-related metabolic processes, as well as other metabolic pathways like butanoate metabolism, the metabolism of antioxidants (such as ascorbate and aldarate), the one-carbon pool by folate and peroxisomal-related pathways. On the other hand, down-regulated proteins did not yield significant results in GO term enrichment analysis but did reveal over-represented KEGG pathways. For instance, the molecular pathway associated with oocyte meiosis stood out, involving the alteration of the abundance of four subunits of serine/threonine-protein phosphatase and peptidyl-prolyl cis-trans isomerase. The significantly up- and down-regulated proteins with effect sizes of order greater than 2.5, potential biomarkers of the DMSO cytotoxic effects, are listed in **Figure 5A**.

**Figure 5.**
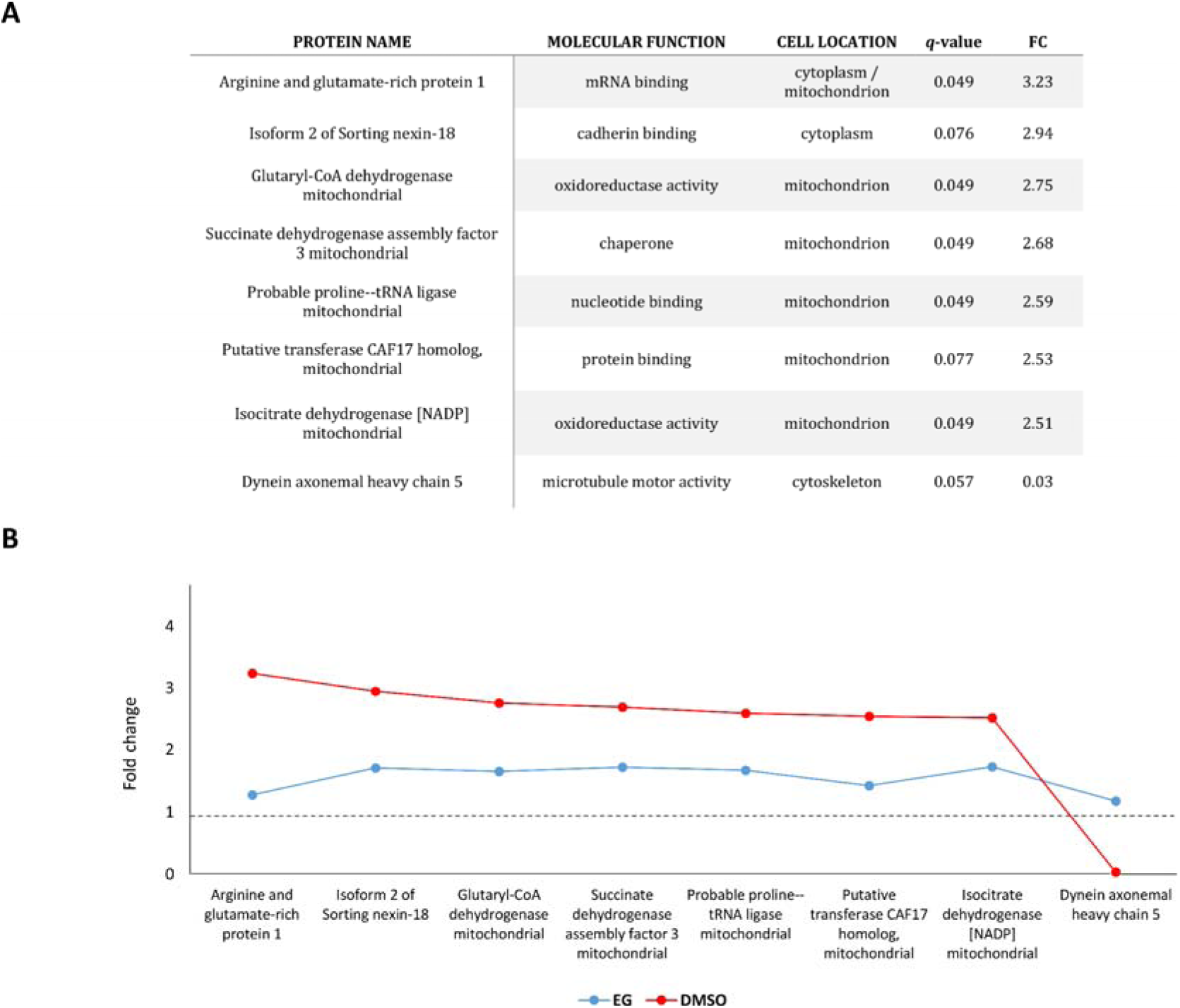
Modulation of Proteins by DMSO and EG Treatments. **A)** List of DMSO-modulated proteins (t-test, q < 0.1) with effect sizes higher than 2.5 or lower than 0.4 for up- or down-regulation, respectively. This list includes their annotation based on the Swiss-Prot database description (“Protein name”), their molecular function, cellular location, significance (*q*-value), and fold-change (FC), after excluding proteins annotated as of bacterial origin. **B)** Fold-changes of the proteins listed in panel A obtained with the DMSO and EG treatments. In all cases, the trends remain consistent between both cryoprotectants, with less extreme FC values with EG compared to DMSO, except for dynein-HC 5, which exhibits similar relative abundance to controls with EG (FC = 1.17) while being significantly down-regulated with DMSO (FC = 0.03).

Contrastingly, a more confined set of proteins experienced significant changes in the protein abundance due to EG treatment, with 88 displaying up-regulation and 13 down-regulation. This widespread protein up-regulation (p < 0.001 in a binomial test versus a 1:1 ratio for up:down regulated proteins) follows a similar trend to the effects detected with DMSO. In contrast to the broader impact of DMSO, EG induces fewer protein modulations, without contributing additional insights beyond those discussed in section 3.3.1. It is worth noting that when examining the relative abundance trend for proteins listed in **Figure 5A**, regardless of statistical significance for EG samples, except for the dynein-HC 5, all FC trends remain consistent between samples treated with any of the two CPAs. In all cases, the effect size (FC) values are substantially lower in EG compared to DMSO-treated samples, as illustrated in **Figure 5B**.

### 3.4. Proteome alterations induced by the monitored slow freezing

No modulated proteins were detected in the oocyte samples treated with EG followed by monitored slow freezing (EG-MSF) when compared to those treated only with EG followed by flash freezing in N_2_(l). In the DMSO-MSF treatment, significant alterations were detected in only eleven proteins, with only seven of these providing informative functional information (**File S1-wsM**). Of these seven, the three up-regulated proteins (FC > 1.5) are expressed in lysosomes and play roles in lipid degradation (phospholipase B-like 1), oxidative stress (lysosomal Pro-X carboxypeptidase) and apoptosis (cathepsin L1). However, it is important to note that the abundance of all three of these lysosomal proteins was already increased with the DMSO treatment, and it is further increased after de MSF treatment.

## 4. DISCUSSION

In this study, we investigated the effects of the two most commonly used cryoprotectants (CPAs), dimethyl sulfoxide (DMSO) and ethylene glycol (EG), along with the process of monitored slow freezing (MSF) followed by thawing, on the oocytes of *Mytilus galloprovincialis*, a valuable marine bivalve species for the aquaculture industry. Our comprehensive analysis encompassed functional, structural, and molecular data. This integrative approach provides valuable insights into optimising oocyte cryopreservation protocols useful in conservation and reproductive biotechnology, shedding light on the molecular mechanisms underlying the limited success reported to date in cryopreserving oocytes from marine invertebrate species.

Our findings provide evidence that neither DMSO (1.5 M) nor EG (1.5 M) treatments induced noticeable alterations in the ultrastructure of the oocyte, particularly in the vitelline membrane (VM), a critical structure directly involved in the sperm-oocyte recognition during fertilisation. Therefore, we did not detect any structural CPA-induced structural damage, even though DMSO is well known for altering cell membrane dynamics and inducing pore formation [65]. These results are in line with those obtained in sea urchin (*Paracentrotus lividus*) oocytes treated with the same CPAs at 0.5 M, with detrimental effects observed at a higher concentration (3 M) [9]. Additionally, there was no significant impact of both CPAs on fertilisation rates compared to those obtained from control oocytes. This is consistent with findings in Pacific oyster (*Crassostrea gigas*), where oocytes treated with 1.8 M EG for 20 minutes also showed no reduction in fertilisation rate [1]. However, in a similar study involving *M. galloprovincialis* oocytes treated with either DMSO or EG at concentrations of 1 M and 2 M for 15 minutes, a significant decrease in fertilisation rate was reported, with a 20% decrease at 1 M and a 40% decrease at 2 M for both CPAs [62]. A similar finding was reported in the marine mussel *Perna canaliculus*, where an average decline of 22 % in fertilisation rate was observed after treating the oocytes with 1.8 M EG for 15 minutes [23].

Additionally, we provide, for the first time, evidence of a significant decrease in D-larvae development rates 72 hours’ post-fertilisation of oocyte samples treated with both CPAs, compared to controls. This decrease was more pronounced with DMSO, with a 41% reduction compared to control, whereas in the EG-treated samples the reduction was 26%. This is in line with the detection of a higher number of features at earlier development stages, *i.e.*, between fertilised eggs and the D-larvae, and suggests that larval development slows down during early stages due to exposure to CPAs. However, whether this effect implies lower viability beyond D-larvae stage remains unclear. This result contrasts with previous findings from similar studies in *P. canaliculus* and *C. gigas*, where no significant decrease in larval development was detected [1,23]. However, they align with results obtained in sea urchin *P. lividus*, where the application of 1.5 M EG for 30 minutes reduced the normal development rate of pluteus larvae by 54 % [49]. To the best of our knowledge, no similar studies have been reported in *M. galloprovincialis*. In contrast, other similar studies have exclusively assessed development rates by using cryopreserved oocytes after thawing [33,62,74].

Despite the absence of any noticeable modification in the oocyte ultrastructure or fertilisation rates, the preceding reported findings of lower development rates are consistent with substantial effects of CPAs at the molecular level. We found evidence that DMSO induced the most significant alterations in the oocyte proteome, modulating the abundance level of 26% of identified proteins, followed by the EG with 16%. Notably, in 99% of the proteins modulated by both CPAs, effect sizes were larger with DMSO than with EG. Taken together, the results from functional and molecular analyses of the analysed oocytes strongly indicate that DMSO is the most cytotoxic CPA tested in our study, consistent with similar findings in previous research [62].

The findings revealing DMSO’s pronounced cytotoxicity compared to EG are further elucidated by analysing enriched molecular pathways and functions within the set of modulated proteins. While these molecular pathways and functions are not directly associated with fertilisation, as evidenced by the unaffected fertilisation rates observed in oocyte samples exposed to both CPAs, they are consistent with a cellular response to CPAs’ cytotoxicity, potentially disrupting some of the mechanisms involved in early development. This could be attributed to changes in the abundance of proteins within the oocytes, primarily encoded by the maternal genome (referred to as maternal factors). These proteins directly influence the phenotype of the oocyte, including its capacity to develop optimally during the early embryonic stage after fertilisation [66]. In fact, some studies on maternal effects in marine organisms suggest that changes in offspring phenotype are often explained by adaptive maternal responses to environmental changes [37]. These findings emphasise the intricate relationship between maternal factors, environmental influences, and offspring development. An important consideration is that changes in the levels of certain proteins in the oocyte, although lacking an obvious link to early development, could play a significant role in meiotic restart and the initial mitotic divisions during the early stages of development after fertilisation. A good example is proteins primarily involved in stress response. Previous findings suggest that these proteins may contribute to the decreased developmental rates observed in *P. canaliculus* when there is an increase in reactive oxygen species (ROS) in oocytes treated with CPAs [23]. This hypothesis gains support from research indicating that mitochondrial ROS act as a limiting factor in the activation of key genes controlling early development in sea urchins [12]. These observations align with our findings, as the exposure to either of the two CPAs resulted in the alteration of proteins associated with energy metabolism pathways, primarily exhibiting mitochondrial oxidoreductase activity (see **Figure 6**).

**Figure 6.**
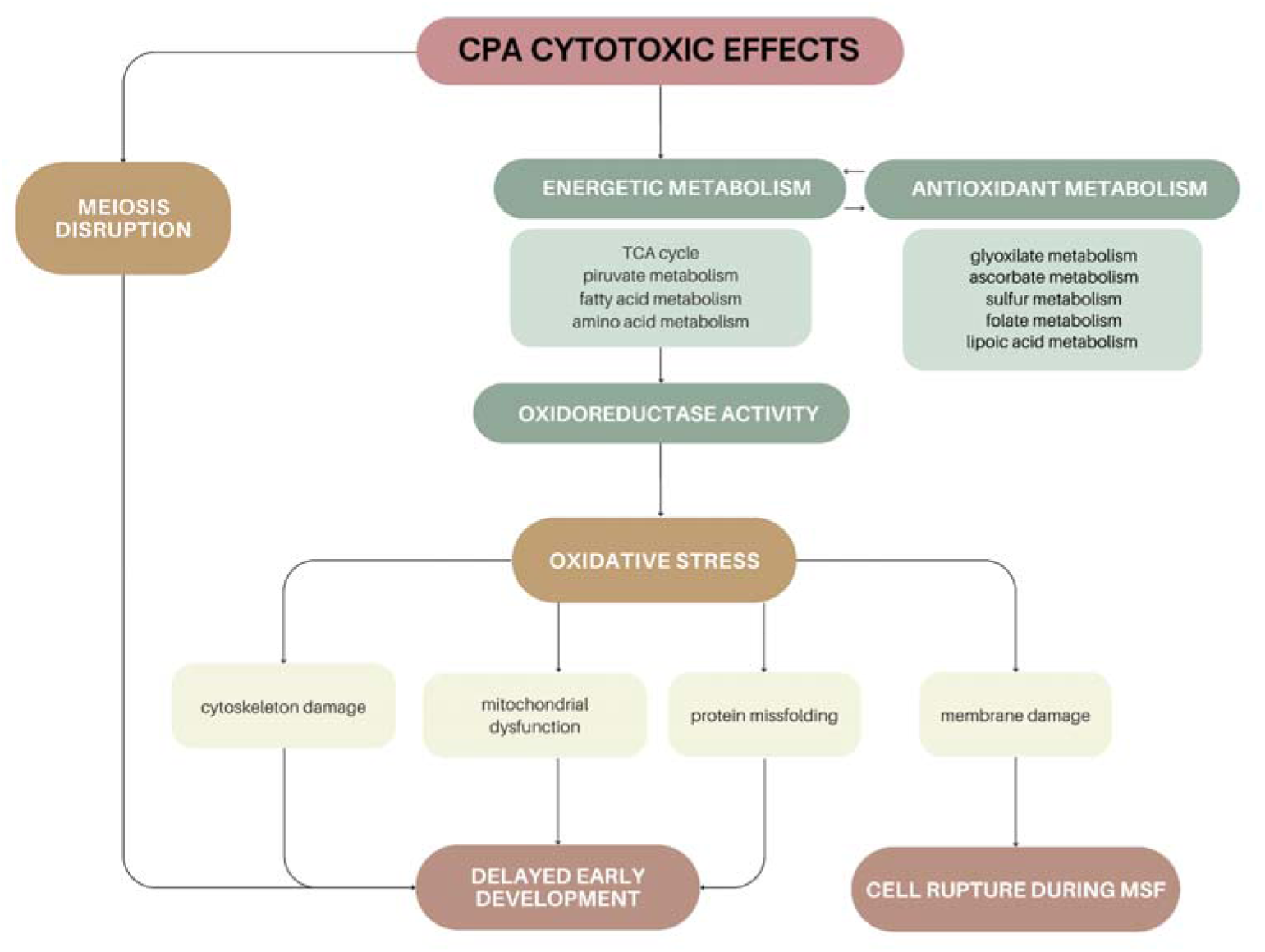
Cellular Damage and Response Network Induced by CPA Cytotoxicity. Illustrative diagram outlining the cellular damage and response network induced by the cytotoxic effects of analysed cryoprotectant agents (CPAs) based on molecular, functional, and ultrastructural analysis at the oocyte level. Predominantly up-regulated pathways correspond to energy metabolism, leading to the subsequent activation of the mitochondrial oxidoreductase activity. This increase in oxidative activity results in intracellular ROS accumulation, triggering oxidative stress. In response, there is an up-regulation of the secondary antioxidant metabolism. However, this cellular response is insufficient, as oxidative stress-induced damages such as cytoskeletal instability, mitochondrial dysfunction, and protein denaturation, along with damages to proteins directly involved in meiosis, slow early development to the D-larvae stage. Additionally, oxidative damage causes membrane impairments, compromising its adaptability to monitored slow freezing (MSF), and culminating in verified cell rupture observed through scanning electron microscopy.

The up-regulation of proteins associated with energy metabolism pathways in oocytes treated with CPAs suggests an elevated metabolic rate linked to mitochondrial activity that may lead to higher ROS accumulation and subsequent oxidative stress [27,55,68]. Such stress may damage mitochondria and the cytoskeleton while impairing protein denaturation mechanisms [45,53]. This is consistent with the detection of up-regulated proteins involved in the cellular response to damage caused by oxidative stress, protecting mitochondria, mitigating ROS production to maintain the right redox balance, and the unfolded protein response (UPR). Specifically, these include the ClpP protease, the SOD Fe/Mn, and the succinate dehydrogenase assembly factor 3, that have been linked to maintaining mitochondrial quality by removing damaged mitochondrial proteins, detoxifying superoxide radicals, and protecting succinate dehydrogenases from the detrimental effects of oxidants, respectively [7,43,44,48]. Also, the observed up-regulation of heat sock proteins (HSP) 10, 60, 70, and 75, proteins involved in UPR, with HSP 70 being directly linked to oxidative stress response [72]. Additionally, it is noteworthy to highlight the detected up-regulation of enzymes associated with glyoxylate metabolism in peroxisomes, organelles involved in free radical scavenging and thus playing an important role in responding to oxidative stress [14]. Similarly, the detection of up-regulated proteins linked to lipoic acid metabolism, with its reduced form known to act as an antioxidant by reacting with ROS [46], supports this line of evidence.

Moreover, the impact of DMSO extends beyond mitochondrial function and the oxidative stress response, affecting secondary metabolic pathways. These include alterations in ascorbate, sulphur, and folate metabolism, which are essential for antioxidant defence and redox homeostasis [21,41,56]. DMSO also induces damage to cytoskeleton-associated proteins such as dynein and those involved in the restart of oocyte meiosis. This aligns with previous reports describing DMSO as a disruptor of cytokinesis by inhibiting cortical reorganisation and polarisation in mouse oocytes [73], providing new potential candidates, along with oxidative stress, to slow down early development.

While CPAs affect the proteome and early embryo development without impacting the oocyte ultrastructure or fertilisation rates, the subsequent monitored slow freezing (MSF)-thawing process resulted in severe structural damage, leading to the complete absence of fertilisation events. These effects included alterations in the vitelline membrane and even leakage of the cellular content of cryopreserved oocytes. Therefore, the freezing-thawing process primarily had negative effects at the oocytés structural integrity, with a lesser impact observed at the molecular level. Specifically, only three up-regulated lysosomal proteins associated with lipolysis, oxidative damage, and apoptosis were detected in the DMSO-MSF samples compared to their respective control oocytes (i.e., those treated with DMSO but not subjected to MSF). However, these proteins were already up-regulated in samples treated only with the CPA, suggesting an extension of the DMSO cytotoxic effects during the MSF-thawing process. Besides the cellular damage caused directly by ice crystal formation during the freezing process, an increase in oxidative stress prior to freezing may compromise the structural integrity of oocytes following the cooling procedure, with high levels of ROS affecting the fluidity and function of the cell membrane [23]. This once again highlights the trade-off between cryoprotection and cytotoxicity. It remains unclear whether the rupture is caused by the ineffective cryoprotective effect of CPAs at the tested time and concentration, failing to mitigate ice crystal formation, or by their cytotoxic effects via oxidative stress, compromising the oocyte’s ability to tolerate low temperatures.

Our results align with the low and highly inconsistent success rates reported in the cryopreservation of marine invertebrate oocytes. For instance, cryopreserved oocytes in *P. canaliculus* mussels have shown null post-thawing fertilisation rates using either DMSO or EG [2]. A similar result was observed in *M. galloprovincialis* when using DMSO, with less than 5% of oocytes fertilised [62]. Comparable results have been reported in oysters, where fertilisation rates are often low or highly inconsistent, with some reports claiming success rates from 0% to 49%, depending on the cryopreservation conditions used [1,60]. Better fertilisation rates were reported in *P. canaliculus* and *M. galloprovincialis* when EG was supplemented with trehalose, a membrane protector, or Ficoll, an osmotic stress-reducing agent [2,33,62]. However, in oysters, EG supplemented with trehalose did not lead to improved outcomes [60]. Despite the reported improvement in fertilisation rates in mussels, this did not translate into successful development beyond the early stages. The D-larvae development rates from cryopreserved oocytes using EG supplemented with trehalose or Ficoll were below 1% and 14%, respectively [2,33,62]. This may be attributed to the inability of trehalose and Ficoll to mitigate the toxic effects of EG and DMSO at the molecular level, as reported in the present study. However, recent findings suggest that incorporating antioxidants into CPA solutions could further enhance marine mussel oocyte cryopreservation [74], though the molecular mechanisms underlying these improvements remain unexplored. Our study addresses this gap by providing molecular insights that align with and help explain these reported functional benefits, potentially increasing success rates if CPA cocktails were supplemented with antioxidants such as α-tocopherol. The similarly low success rate observed in fish oocyte cryopreservation, which reaches at most 30% (reviewed in [17]), may also be influenced by the cytotoxic effects of CPAs. Indeed, DMSO has been shown to compromise oocyte membrane integrity in zebrafish [25,51]. The evidence of oxidative damage caused by CPAs in mussel oocytes presented in this study may also be relevant to improving fish cryopreservation protocols, offering potential pathways for enhancing their success rates.

In summary, this study provides new insights into the structural, functional, and molecular effects of the two most commonly used cryoprotectant agents, DMSO and EG, in the cryopreservation of *Mytilus galloprovincialis* oocytes. By comprehensively analysing these effects, we shed light on the persistent challenges limiting the success of oocyte cryopreservation in marine mussels and other invertebrates, contributing to the development of improved protocols. Our findings reveal significant cytotoxic effects at the proteome level, with DMSO causing more pronounced alterations than EG. These effects are primarily linked to increased oxidative stress and disruptions in meiotic-related proteins, suggesting that optimizing cryopreservation strategies should focus on minimizing cytotoxicity, with EG emerging as the preferable option. While these alterations do not initially compromise oocyte ultrastructure or fertilisation rates, they lead to a significant delay in early development and may impair the membrane ability to adapt to subsequent freezing-thawing processes, ultimately resulting in the observed ultrastructural and functional alterations in post-thawed cryopreserved oocytes. Our findings reinforce the strategy of supplementing CPAs with antioxidants in marine mussel oocytes—and potentially in other marine invertebrate species—to effectively reduce cytotoxic effects, thereby improving the viability of cryopreserved oocytes. Furthermore, they strongly suggest that future research should focus on developing cryoprotectant cocktails that combine multiple compounds to minimize toxicity while preserving cryoprotective efficiency.

## AUTHORS’ CONTRIBUTION

**Sofía Blanco**: Conceptualization; methodology; investigation; formal analysis; writing—original draft, review and editing. **Sara Campos**: Methodology; investigation; writing—review and editing. **Patricia Reboreda**: Methodology; investigation; writing—review and editing. **Estefanía Paredes**: Conceptualization; supervision; methodology; investigation; writing—review and editing. **Angel P. Diz**: Conceptualization; funding acquisition; supervision; methodology; investigation; formal analysis; writing—original draft, review and editing.

## DATA AVAILABILITY

The mass spectrometry proteomics data have been deposited to the ProteomeXchange Consortium via the PRIDE partner repository with the dataset identifier PXD053176 and 10.6019/PXD053176.

## ACKNOWLEDGEMENTS

Manuel Marcos for performing the LC–MS/MS analyses and Inés Pazos for performing the scanning electron microscopy (SEM) analyses at the Structural Determination, Proteomics and Genomics unit and the Electron Microscopy unit of CACTI (Universidad de Vigo), respectively.

